# Leveraging Pretrained Vision Transformers for classifying Alcohol Use Disorder using Raw Resting-State EEG

**DOI:** 10.64898/2026.01.14.699473

**Authors:** A. Bingly, C.D. Richard, B. Porjesz, J. L. Meyers, D.B. Chorlian, C. Kamarajan, G. Pandey, W. Kuang, A. K. Pandey, S. Kinreich

## Abstract

Alcohol Use Disorder (AUD) is a prevalent and debilitating neuropsychiatric condition characterized by compulsive alcohol consumption, impaired control, and negative emotional states, affecting about 28 million adults in the United States. Despite its significant public health burden, there are few objective biomarkers and no reliable neurophysiological tools to assist in its clinical diagnosis. In this study, we investigated the potential of deep learning to classify individuals with AUD using raw resting-state electroencephalogram (EEG) data. EEG recordings were obtained from the Collaborative Study on the Genetics of Alcoholism (COGA), a large, longitudinal, multi-site dataset. The initial cohort included a total of 5,402 recordings from 2,710 participants (aged 12–83, mean age 24; 1,338 males and 1,372 females). To reduce confounding factors, we applied demographic matching, and to address class imbalance, we applied undersampling. Minimal preprocessing was applied to preserve the raw EEG features. We utilized EEGViT, a hybrid deep learning architecture that combines convolutional patch embedding with a Vision Transformer (ViT) pretrained on ImageNet, thereby enabling end-to-end learning directly from raw EEG input. The analysis was stratified by sex and age, and all groups were age-matched. To validate the generalization of the model, models were also trained for Cannabis Use Disorder (CUD) and Opioid Use Disorder (OUD). Results for the AUD model showed a classification accuracy of approximately 56% in the overall dataset, 54% for males, and 58% for females. The CUD model showed an accuracy of about 63% with 59% for females and 69% for males. The OUD model showed an accuracy of about 63% with 61% for females and 65% for males. Temporal analysis indicated that the model’s performance varied across time intervals, with higher accuracy observed in later minutes compared to earlier ones. While modest, these findings underscore the potential of transformer-based models in psychiatric classification using raw EEG data and provide a foundation for future development of EEG-based diagnostic tools for AUD.

## 1 Introduction

Alcohol Use Disorder (AUD) is a common and debilitating mental health condition marked by excessive habitual drinking and loss of control over drinking.[1] According to the 2024 National Survey on Drug Use and Health, 27.9 million people ages 12 and older had AUD in the past year in the United States [2]. Moreover, estimates suggest that alcohol played a role in at least 7.1% of emergency department visits [3], and an analysis of death certificates showed that deaths involving alcohol among people ages 16 and older accounted for 99,017 [4]. These alarming facts highlight the critical importance of identifying objective and accurate biomarkers predictive of AUD diagnosis and outcomes. Such insights could contribute to preventive strategies and treatments, potentially intervening before the addiction fully develops.

Over the past several decades, AUD neuroimaging research using magnetic resonance imaging (MRI)[5] [6], functional MRI (fMRI)[7], and electroencephalogram (EEG) [8] [9] [10] recordings has focused on developing predictive models and identifying biomarkers associated with the disorder. Studies using structural MRI have found cortical thinning and volume reductions in frontal and temporal regions associated with AUD, which have been used as input features for machine learning (ML) algorithms such as support vector machines (SVM) and random forests to create significant AUD classifiers [5] [6]. fMRI has been used to detect altered brain connectivity patterns, particularly in the default mode and reward networks [7]. EEG-based approaches using features such as spectral power values or event-related potentials (ERPs) have produced significant models that predict people with AUD[8]. Despite significant advances, the ability to reliably detect individuals with AUD or accurately predict vulnerability based on biomarkers remains limited, largely due to the absence of objective, scalable diagnostic tools and the heterogeneity of clinical presentations and brain development [11, 12]. Moreover, while existing studies show encouraging results, they are often constrained by small sample sizes[13], inconsistent data processing pipelines, and a lack of external validation, all of which hinder generalizability across populations [14] [15].

Our current work focuses on classifying AUD individuals using a substantial amount of EEG data from the Collaborative Study on the Genetics of Alcoholism (COGA). COGA is a landmark, longitudinal, multi-site research project aimed at identifying genetic, neurobiological, and environmental factors that contribute to the development and persistence of AUD and related conditions.[16] EEG offers a noninvasive, cost-effective method for capturing brain activity with high temporal resolution and has shown promise in identifying neurophysiological markers of AUD [17]. Among the various neuroimaging paradigms, resting-state assessment is particularly attractive for clinical applications due to its portability, simplicity, minimal task demands, reproducibility, and consistency across imaging and electrophysiological modalities. [18]

Resting state EEG findings, including elevated theta power at rest, most notably in the frontal, central, and parietal regions, have been identified among AUD relapsers[19]. Resting parietal-occipital alpha power is consistently found to be reduced in AUD, while increased frontal-central beta activity is found in both active and abstinent individuals [20]. Resting-state EEG functional connectivity shows enhanced slow-wave coupling (delta, theta, alpha) and stronger beta-band correlations within the parietal region [21].

While traditional methods for classifying AUD and identifying AUD biomarkers have relied on handcrafted features input to models such as logistic regression and k-nearest neighbors, more recent efforts have used convolutional and recurrent neural networks to classify AUD directly from raw or minimally processed EEG [9] [10]. Overprocessing of the signal may obscure subtle, yet important, discriminative features and introduce bias, thereby reducing the model’s ability to learn meaningful patterns [22]. Preserving the raw signal enables the model to learn potentially informative features that may be distorted or lost through aggressive preprocessing. This is particularly important in the context of AUD, where the question of widely accepted EEG biomarkers is still open.

Therefore, in the current study, we implemented an advanced deep learning model trained on the COGA EEG resting-state dataset. Deep learning models, particularly convolutional and recurrent neural networks, have demonstrated strong capabilities in learning directly from raw physiological signals, including EEG, without the need for extensive feature extraction [23] [24]. [24]. More recently, transformer-based architectures, originally developed for natural language processing [25], have shown substantial promise in modeling EEG data due to their ability to capture long-range dependencies and flexible attention mechanisms [26] [27]. Unlike Convolutional Neural Network (CNNs) or Recurrent Neural Networks (RNNs), transformers do not rely on fixed kernel sizes or sequential processing, making them well-suited to handle the non-stationary and multiscale characteristics of EEG signals. Specifically, variants such as Vision Transformers (ViT) have been successfully adapted for time series and physiological data, and recent work has demonstrated their competitive performance in EEG-based classification tasks [28] [29], including cognitive workload assessment [30], sleep staging [31], and seizure detection [32].

Moreover, recent studies have shown that leveraging pretrained transformer models, either on large image or EEG-specific datasets, can provide beneficial inductive biases that improve classification performance when fine-tuned on smaller datasets [28] [33]. A recent review further highlights transfer learning with pretrained transformers as a promising strategy in EEG analysis [26]. To further assess the robustness of the transformer model, we evaluated its performance on other substance use disorders in the COGA dataset, including Cannabis Use Disorder (CUD) and Opioid Use Disorder (OUD).

This study aims to classify AUD individuals by leveraging a transformer-based deep learning architecture specifically adapted for raw EEG and by using a large, longitudinal dataset. We will also evaluate the models with CUD and OUD data. Moreover, we aimed to investigate whether raw EEG signals contain discriminative features that could inform future development of automated, EEG-based screening tools for AUD.

## 2 Method

### 2.1 Dataset

Our cohort includes individuals from the COGA database. Initiated in 1989, COGA is funded by the National Institute on Alcohol Abuse and Alcoholism (NIAAA) and currently spans ten research centers across the United States [34]. The COGA study recruited nearly 18,000 individuals from families with a high density of AUD. Participants underwent behavioral, clinical, and neurophysiological evaluation, assessed repeatedly across the lifespan, providing extensive longitudinal data, including EEG recordings [16]. All participants provided informed consent, and data collection protocols have been approved by Institutional Review Boards (IRBs) at each participating site.

### 2.2 Participants

A total of 2710 individuals were included in the analysis, comprising 1,338 males and 1,372 females. Participant ages ranged from 13 to 83 years, with a mean age of 24 years. The data included multiple participants’ visits, ranging from 1 to 9, with a mean of 2 visits per participant. At each visit, participants were assessed using the Semi-Structured Assessment for the Genetics of Alcoholism (SSAGA), which is a diagnostic instrument for AUD and other substance use disorders, in addition to EEG, neuropsychological, and behavioral assessments. Each participant was diagnosed as either having AUD (case group) or not having AUD (unaffected group), based on criteria outlined in the *Diagnostic and Statistical Manual of Mental Disorders, Fifth Edition (DSM-5)* [35]. In total, the dataset included 5402 visits, of which 2701 were labeled as AUD and 2701 were labeled as non-AUD (Table 1)

### 2.3 EEG Acquisition

Resting-state EEG closed eyes data were acquired using a 64-channel EEG electrode cap, following the International 10-20 system for electrode placement. Recordings were acquired at a sampling rate of 500 Hz. During the recording, participants were seated in a comfortable chair in a dimly lit, acoustically and radio-electrically shielded room. They were instructed to keep their eyes closed and remain relaxed to minimize motor artifacts.

### 2.4 Preprocessing Pipeline

The EEG preprocessing pipeline is illustrated in Fig. 1, and each step is detailed in the following subsections.

**Figure 1.**
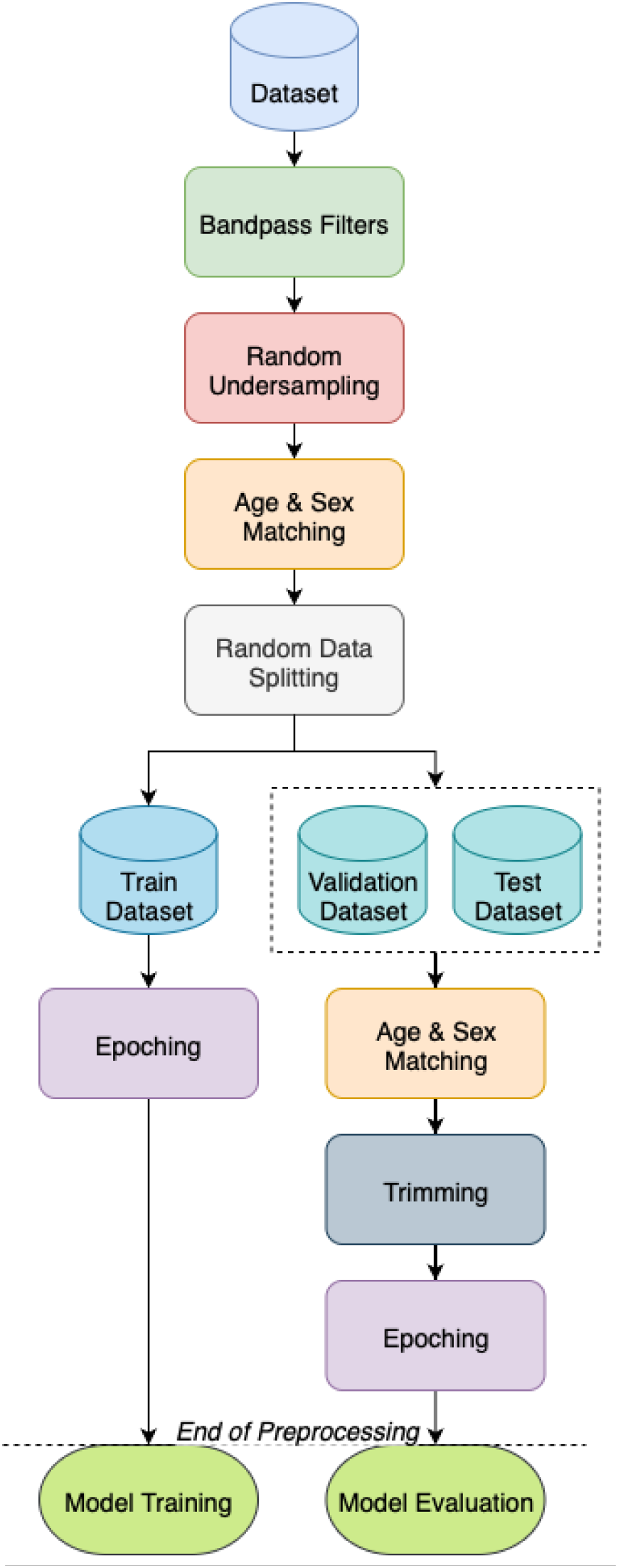
The preprocessing pipeline for EEG data.

#### 2.4.1 Bandpass Filters

To enhance signal quality and denoise signal artifacts due to motor, drift, and electrical interference, the EEG recordings were bandpass filtered between 1Hz and 60Hz [36, 37]. Filtering was performed using a zero-phase finite impulse response (FIR) filter, commonly recommended for EEG preprocessing due to its ability to eliminate phase distortion while preserving the temporal structure of the signal [38, 39]. Uniform filtering was applied across all channels to maintain data integrity and consistency for downstream analyses.

#### 2.4.2 Random Under-sampling

Given the inherent class imbalance in the dataset, where the number of unaffected participants significantly exceeds the number of cases, it was crucial to address this disparity before model training. Class imbalance can lead to biased model performance, typically favoring the majority class and resulting in poor generalization to the minority class [40]. To mitigate this, we applied random under-sampling to the majority class (unaffected), resulting in a more balanced distribution, a strategy proven effective in the physiological signal classification context [41].

#### 2.4.3 Demographic Matching

With the influence of age on both brain structure and function, along with evidence of sex-related differences in brain activity [42][43], we performed age and sex matching between the cases and the unaffected [21]. Each AUD case was matched with an unaffected of the same sex and within a ±5-years age range, consistent with our prior work and chosen to optimize the trade-off between demographic parity and sample retention [21].

#### 2.4.4 Validation and Testing

The same preprocessing pipeline was applied to the validation and test sets, with additional constraints to ensure robust evaluation. The constraints include strictly age- and sex-matched. All recordings were trimmed to a standardized 4-minute duration. Moreover, it was ensured that no more than 1 recording per participant was included. These measures and precautions minimized the risk of confounding factors influencing evaluation outcomes, including demographic and sampling variability.

#### 2.4.5 Epoching

Each EEG signal was divided into 1-second non-overlapping epochs, yielding multiple samples of 500 time points per channel [28], as illustrated in Fig. 2A. This window size was chosen to balance temporal resolution and computational feasibility. Each epoch was treated as an independent sample during training and evaluation.

**Figure 2.**
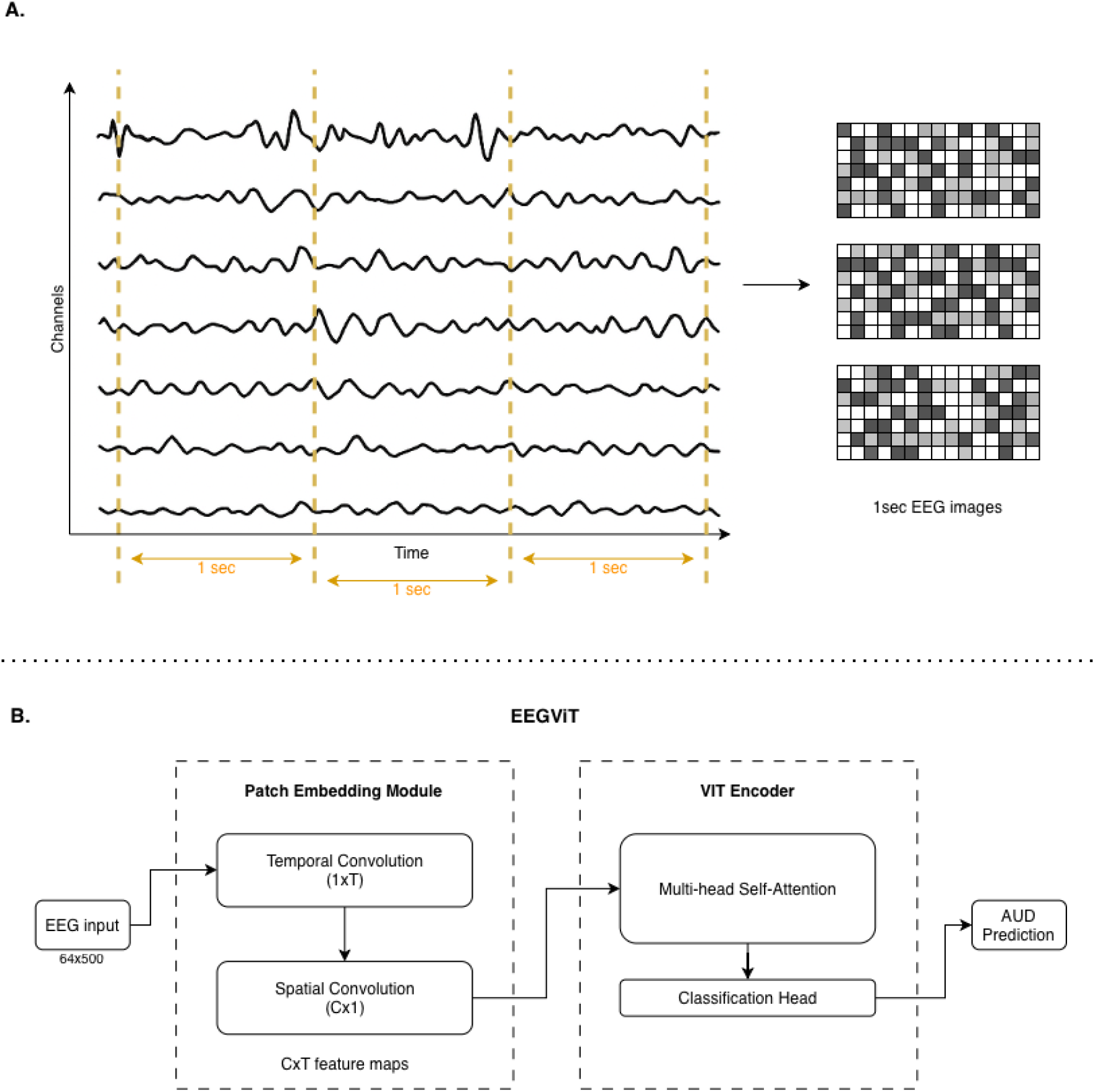
Epoching and the EEGViT model architecture. A. The EEG signal is divided into 1-second non-overlapping epochs. These one-second epochs serve as input to the model as images. B. The model consists of a patch embedding module that applies temporal and spatial convolutions to the raw EEG signal, followed by a Vision Transformer (ViT) encoder pretrained on ImageNet. The output is then passed through a classification head to predict AUD.

### 2.5. Model - EEGViT

We employed a hybrid deep learning model called EEGViT, adapted from the paper, *ViT2EEG, Yang and Modesitt (2023)* [28]. It is based on the Vision Transformer (ViT) architecture [44], with modifications to better capture the spatiotemporal structure of raw EEG signals. Specifically, the model incorporates a two-step convolutional embedding process prior to the transformer layers. The model operated on 1-second EEG segments, a matrix of 64×500 (channels x time). These short segments were used due to the computational and sequence length limitations of the ViT model, which make it infeasible to process entire 4-minute recordings directly. Segmenting the EEG in this way also standardizes input size and enables batch-wise training.

The patch embedding module consisted of two convolutional layers designed to mimic temporal and spatial filtering typically used in EEG signal processing. The first layer applies a 1×T convolution across the temporal dimension for each channel. This temporal convolution can be interpreted as a learnable set of band-pass filters, allowing the model to extract frequency-specific features directly from the raw signal [45]. The output feature maps capture relevant temporal dynamics associated with the EEG signal.

The second layer applies a depth-wise C×1 convolution across the spatial (channel) dimension, acting as a frequency-specific spatial filter. This enables the model to learn localized spatial features by aggregating information across EEG channels, similar to spatial filtering techniques such as common spatial patterns (CSP), but in a data-driven manner.

The output of the patch embedding module is a set of C×T feature maps, which are then reshaped into fixed-size patches and passed to a standard ViT encoder. Fig. 2B illustrates the model architecture.

The core of the model is a ViT encoder pretrained on ImageNet. Although ImageNet pretraining is originally intended for visual tasks, recent work has shown that pretrained ViT backbones provide useful inductive biases and generalizable priors that benefit EEG-based classification tasks, even in domains with significant differences from natural images [46] [47]. The encoder employs multi-head self-attention mechanisms to model long-range dependencies across the temporal and spatial dimensions of the EEG data. The encoded representations are passed through a final classification head to predict AUD.

### 2.6 Training Procedure

The EEGViT model was trained using a combination of optimization techniques, fine-tuning strategies, regularization methods, and data augmentation to enhance generalization and performance. The AdamW optimizer was used, decoupling weight decay from gradient updates, allowing for more stable training and improved regularization [48]. The learning rate followed a cosine annealing schedule, enabling gradual decay and periodic resets to help escape local minima [49]. The initial learning rate was set to 1e-4, and warm restarts were applied every 5 epochs.

#### 2.6.1 Progressive Fine-Tuning Schedule

During training, a progressive fine-tuning schedule was employed to gradually adapt different parts of the model. Initially, only the patch embedding layers were trained for the first two epochs to stabilize early learning. Next, the classifier and positional embeddings were unfrozen and trained with a moderately high learning rate of 1e-4 and weight decay of 1e-4, combined with a cosine annealing scheduler over 5 epochs. Subsequently, the top transformer encoder layers (layers 9, 10, and 11) and layer normalization parameters were unfrozen and fine-tuned with a reduced learning rate of 1e-5, also following a cosine annealing schedule. The remaining transformer encoder layers were progressively unfrozen in stages across subsequent epochs until the entire transformer backbone was trained. This gradual unfreezing approach allowed for stable and effective transfer learning, mitigating sudden shocks to pretrained weights and improving convergence [50]. The transition between each phase was based on early stopping, and the early stopping metric was reset in each phase to avoid premature convergence [51].

#### 2.6.2 Data Augmentation

Data augmentation strategies were implemented during training to improve the model’s robustness to variability in the EEG signal. Gaussian noise with a mean and a standard deviation similar to the dataset was added to the input segments to simulate sensor or environmental noise [45]. In addition, the start point of the first 1-second input window was randomly shifted within a 0 to 1-second range, creating variability in the temporal alignment of the segments. These augmentations were applied on-the-fly during training and were not used for validation or testing, ensuring the integrity of model evaluation [52].

### 2.7 Additional Analysis

#### 2.7.1 Time-stratification

To further investigate the temporal dynamics of model performance, we examined how classification accuracy varied across different segments of the EEG recording. Specifically, a subset of the data (participants with 5-minute recordings) was used to assess the model’s accuracy on all possible contiguous time windows within this time span. For example, given a 5-minute EEG recording, the following combinations were assessed: 0-1min, 0-2min, 0-3min, 0-4min, 0-5min, 1-2min, 1-3min, 1-4min, 1-5min, 2-3min…

#### 2.7.2 Model Evaluation with CUD & OUD data

To evaluate whether the proposed EEGViT model captures features that are specific to AUD or generalizable to other substance-related conditions, we extended our analysis to additional use disorders available within the COGA dataset - CUD and OUD. For each disorder, participants were categorized as case or unaffected, where cases met diagnostic criteria for the respective disorder based on DSM-5, similar to AUD. The number of participants in each of the analyses (CUD, OUD) is provided in Tables S1 & S2 in the supplementary file.

The same preprocessing pipeline was applied to the EEG data, including bandpass filtering, demographic matching, and random undersampling to achieve balanced class representation. Each model was trained independently for each disorder using the EEGViT architecture, with identical training procedures and hyperparameter configurations. Training was conducted from scratch for each disorder to ensure that model weights were not influenced by prior AUD-specific learning.

Performance metrics (accuracy, precision, recall, and F1-score) were computed for each disorder using an independent test set. This analysis provided insight into whether shared or disorder-specific EEG signatures could be captured using the same transformer-based framework and whether pretrained ViT representations could generalize across different domains of substance-related neurophysiological alterations.

#### 2.7.3 Age-stratification

EEG characteristics are known to vary substantially with age due to developmental and aging-related changes in brain structure and function. To evaluate whether age-related variability influenced model performance, we conducted a post hoc age-stratified analysis using the same trained EEGViT model. Specifically, we selected a subsample of participants aged 20–40 years, representing a relatively homogeneous adult cohort in which both brain maturation and age-related decline effects are minimal. This range was chosen to reduce potential confounding from developmental (adolescent) or aging (older adult) neural differences that may alter EEG spectral power or connectivity patterns independent of AUD status.

The previously trained EEGViT model was applied directly to this age-restricted test set without further fine-tuning or retraining. The same preprocessing and matching procedures were maintained to ensure consistency. Performance metrics were computed to assess whether the model’s classification ability improved within this narrower age window, providing insight into the influence of age heterogeneity on EEG-based AUD detection.

## 3. Results

### 3.1.1 AUD Model

The EEGViT model was used to classify individuals with AUD versus those who are unaffected. The results presented in Table 2 indicate a moderate classification outcome for the analysis that included the entire test dataset, with an accuracy of 56.02%, a precision of 55.80%, a recall of 86.52%, and an F1-score of 67.84%. The evaluation was also repeated after sex stratification, the model performed with an accuracy of 58.33%, a precision of 57.97%, a recall of 86.96%, and an F1-score of 69.57% on the females; and with an accuracy of 53.66%, a precision of 53.62%, a recall of 86.05%, and an F1-score of 66.07% on males.

**Table 1.** The distribution of the participants for the train, validation, and test datasets.

|  | #Participants<br>(% of total) | #trials<br>(% of total trials) | %AUD | Approx. no.<br>of samples |
| --- | --- | --- | --- | --- |
| **Total** | 2710 | 5402 | 50% | NA |
| **Train** | 2170(80.07%) | 4862(90.00%) | 50% | 305k |
| **Validation** | 270 (9.96%) | 270 (5.00%) | 50% | 64k |
| **Test** | 270 (9.96%) | 270 (5.00%) | 50% | 64k |

**Table 2.** Classification results for the AUD Model.

| Age Group | Sex Group | Accuracy | Precision | Recall | F1 Score |
| --- | --- | --- | --- | --- | --- |
| All | All | 56.02% | 55.80% | 86.52% | 67.84% |
| All | Females | 58.33% | 57.97% | 86.96% | 69.57% |
| All | Males | 53.66% | 53.62% | 86.05% | 66.07% |
| 20-40y | All | 56.76% | 58.70% | 84.38% | 69.23% |
| 20-40y | Females | 56.45% | 56.00% | 84.85% | 67.47% |
| 20-40y | Males | 57.14% | 61.90% | 83.87% | 71.23% |

Fig. 3 illustrates model accuracy for varying combinations of segments of time within the 5-minute resting-state EEG recording. Notably, model performance was lowest in early segments (e.g., 0–1 minute; accuracy = 0.56) and tended to increase in later portions of the recording, peaking between 3–5 minutes with an accuracy of 0.62.

**Figure 3.**
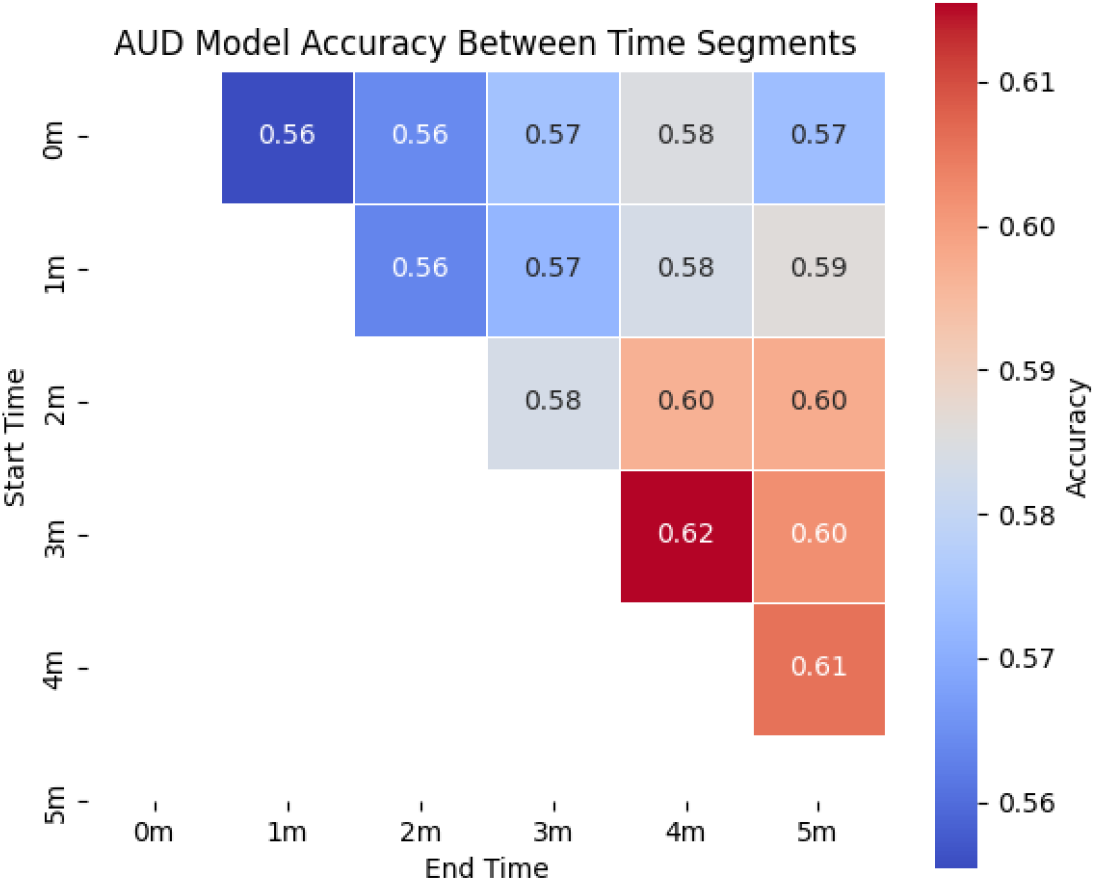
Time-stratification analysis. The figure shows the accuracy of AUD prediction across varying time windows.

### 3.1.2 Model Evaluation with CUD & OUD data

For the CUD model (Table 3), overall classification performance was moderate, achieving an accuracy of 63%, precision of 62%, recall of 88%, and an F1-score of 73% across all participants. When stratified by sex, the model performed better among males (accuracy = 69%, F1 = 80%) compared with females (accuracy = 59%, F1 = 66%).

**Table 3.**
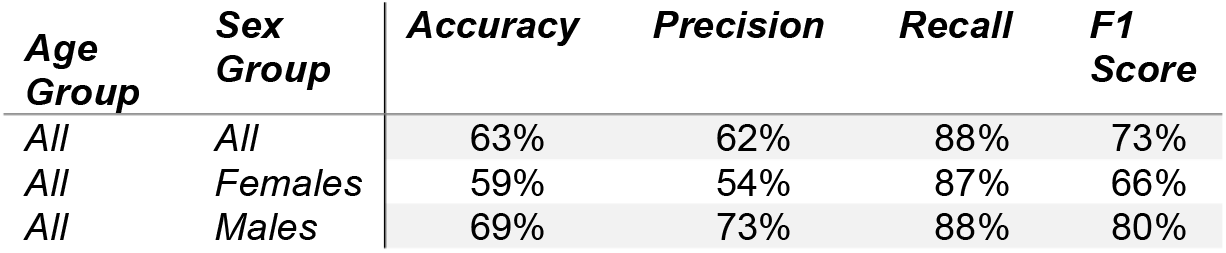
Classification results for the CUD Model.

| Age Group | Sex Group | Accuracy | Precision | Recall | F1 Score |
| --- | --- | --- | --- | --- | --- |
| All | All | 63% | 62% | 88% | 73% |
| All | Females | 59% | 54% | 87% | 66% |
| All | Males | 69% | 73% | 88% | 80% |

For the OUD model (Table 4), the overall performance was similar, with an accuracy of 63%, a precision of 60%, a recall of 76%, and an F1-score of 67%. In sex-specific analyses, females exhibited slightly higher recall (81%) but lower precision (58%), yielding an F1-score of 68%, while males showed more balanced precision (63%) and recall (71%) with an F1-score of 67%.

**Table 4.**
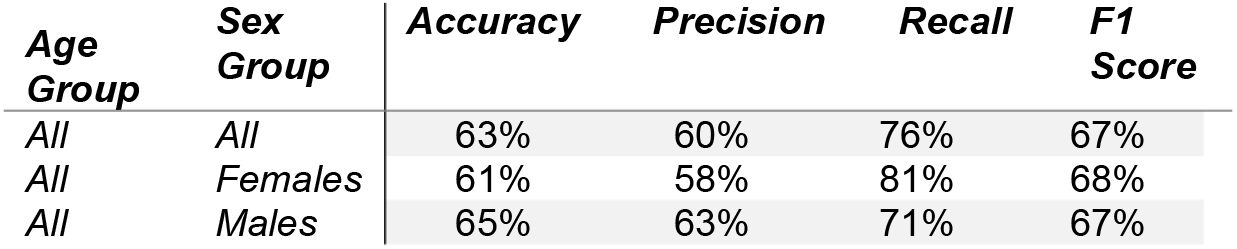
Classification results for the OUD Model.

Both models exhibited consistent performance across evaluation metrics, indicating that the EEGViT framework can be applied to additional substance use disorder classifications using EEG data.

## 4 Discussion

In this study, we explored the feasibility of using a pretrained transformer-based deep learning architecture to classify individuals with AUD from raw resting-state EEG data. Leveraging a large, longitudinal dataset from COGA, our model was trained end-to-end on 1-second raw EEG segments using minimal preprocessing. Our AUD model achieved a modest overall accuracy on the whole sample, and showed similar results when stratified by sex and age. Additionally, the model performance at different time intervals showed higher accuracy in the later minutes compared to the earlier ones. The CUD and OUD models achieved similar results, while the accuracy on the male subset was observed to be slightly higher than that of the female subset. This study provides meaningful insights into the current capabilities and challenges of EEG-based AUD detection using transformer models.

Our results demonstrate valuable classification performance for AUD achieved using a pretrained vision transformer model[53]. The classification results within the narrower age range of 20-40 years were largely consistent with those from the full dataset, suggesting that age heterogeneity did not substantially bias model performance when based on raw data. Nonetheless, the restricted age analysis indicated slightly higher F1-scores in males, aligning with the broader sex-specific trend observed across this study’s models. These findings suggest that while EEGViT model remains robust across adult age ranges, future studies with larger and more evenly distributed age cohorts are needed to better characterize potential age-related differences in EEG-based substance use disorder classification.

Advances in AI and machine learning are creating new venues for classifying and diagnosing individuals with AUD using neuroimaging data. In the current work, we present a first attempt to utilize an EEG-pretrained Vision Transformer model to classify AUD. Using a pre-trained model reduces the reliance on large datasets, a resource often scarce in neuroimaging research[54]. Additionally, pretrained models act as a strong initialization, guiding optimization toward better generalizable solutions and often achieving higher accuracy with less training time and computational cost than models trained from scratch[54]. By capturing rich, transferable representations, such as linguistic patterns in text or visual features in images, pretrained models enable faster convergence, improved robustness to distribution shifts, and strong baseline performance across a variety of domains[55].

The model was trained on raw EEG signals, without relying on handcrafted features. We hypothesized that advanced algorithms could identify patterns that conventional preprocessing might obscure or eliminate. However, the moderate model performance indicates that further refinement of the model or training process is needed to enable effective classification. Interestingly, the results show relatively higher recall, suggesting the model is more sensitive in identifying AUD cases than it is specific in rejecting non-AUD cases. This may have clinical relevance in settings where identifying individuals at risk is prioritized over minimizing false positives[56].

Similar results were observed when the analysis was stratified by sex. The deep learning models were unable to distinguish sex-related variability in EEG signals from differences in the manifestation of AUD-related neurophysiological markers. Prior literature suggested sex-related heterogeneity in both brain structure and the clinical presentation of AUD[57]. For example, EEG beta power is generally higher in young males with AUD, and they usually exhibit greater interhemispheric connectivity, whereas young females with AUD tend towards enhanced interhemispheric connectivity [58]; P3 appears to be particularly sensitive to alcohol misuse, with males exhibiting a lower amplitude than young females [58]. Future analyses will require further model development to improve sensitivity to sex-specific differences.

Beyond AUD, we also evaluated whether the EEGViT architecture could generalize to other substance use disorders. The models trained for CUD and OUD achieved similar overall accuracies. These results suggest that the model captures EEG features that extend beyond alcohol-specific effects, potentially reflecting shared neurophysiological alterations across substance use disorders[59, 60]. Indeed, a recent review of sixty scientific manuscripts concluded that substance use disorders share significant similarities in their genetic architectures and phenotypic profiles[59].

Interestingly, results show that the model was more accurate in classification when given raw EEG resting-state signals from the later part of the recordings. That is, the model performed better without the first few minutes of the EEG resting-state. This temporal trend may reflect initial, settling, or adaptation effects, where early EEG signals capture residual cognitive or physiological noise (e.g., physical movements, eye blinks, or transition into rest) [61, 62] that may obscure AUD-related features. Alternatively, later segments may contain additional information, allowing the model to extract more discriminative knowledge. These findings suggest that temporal selection may influence EEG-based AUD classification, and future works could explore weighting or segment selection strategies to optimize model input.

There are several factors that may contribute to the relatively low classification performance. First, EEG signals are inherently noisy and subject to substantial inter-individual variability [62], which may limit model generalizability. Although we implemented demographic matching and class balancing strategies, other unmeasured confounders, such as comorbid psychiatric conditions, medication use, or environmental influences, may still impact EEG patterns and confound classification[42]. Second, while ViT-based models offer theoretical advantages in modeling long-range dependencies, they may require larger datasets or domain-specific pretraining to achieve their full potential. The pretraining used in this study (on ImageNet) may not transfer optimally to EEG data, which differs substantially from natural images in structure and information content.

Moreover, the use of short (1-second) EEG epochs, although computationally efficient, may limit the amount of contextual information available to the model. Incorporating longer temporal segments or leveraging hierarchical models that integrate across multiple time scales could improve classification performance. Additionally, the model may benefit from multimodal integration, such as incorporating clinical or genetic information from the COGA dataset alongside EEG, to capture a more holistic representation of AUD risk[63].

Our work adds to the current interest in AI, specifically in pretrained transformers classifying neuroimaging data. Specifically, this work demonstrated the potential of transformer-based architectures to learn discriminative features directly from raw EEG for AUD classification. While the current performance may not yet support clinical deployment, this work lays the groundwork for future studies that aim to refine model architecture, integrate auxiliary data, and explore domain-specific pretraining approaches.

## 5 Limitations and Future Direction

While the current study demonstrates the feasibility of transformer-based EEG modeling for substance use disorder classification, several limitations warrant consideration.

Although the COGA dataset is among the largest publicly available EEG resources on AUD, deep learning models would benefit from a more diverse and larger dataset. It would enable more generalizability and also enable the use of more complex model architectures that can leverage both temporal and spatial dependencies within and across EEG channels. Additionally, the use of short, 1-second epochs may restrict the temporal context available to the model. Longer window sizes or models capable of integrating multi-scale temporal dynamics could enhance discrimination of use-related neural patterns. In addition, the ViT architecture used here was pretrained on natural images, which differ markedly from EEG signals. Additionally, explainable AI techniques such as attention-based saliency mapping should be applied to identify which EEG features or brain regions most strongly drive classification. This could not only improve interpretability but also facilitate the discovery of novel electrophysiological biomarkers of substance use risk.

## Acknowledgements

Support for this research came from the National Institute on Alcohol Abuse and Alcoholism (AA029448) and the Collaborative Study on the Genetics of Alcoholism project grant (U10AA008401).

