## Supplemental file for "Leveraging Pretrained Vision Transformers for classifying Alcohol Use Disorder using Raw Resting-State EEG"

### S1.CUD Dataset

*Table S1: The distribution of the CUD participants for the train, validation and test datasets.*

| | **#participants** *(% of total)* | **#trials** *(% of total trials)* | **%CUD** |
|---|---|---|---|
| **Total** | 969 | 1098 | 50% |
| **Train** | 789*(81.42%)* | 918*(83.60%)* | 50% |
| **Validation** | 90 *(9.29%)* | 90 *(8.20%)* | 50% |
| **Test** | 90 *(9.29%)* | 90 *(8.20%)* | 50% |

The CUD dataset included 969 participants, comprising 528 males and 441 females. Participant ages ranged from 14 to 73 years with a mean age of 36 years.

### S2.OUD Dataset

*Table S2: The distribution of the OUD participants for the train, validation and test datasets.*

| | **#Participants** *(% of total)* | **#trials** *(% of total trials)* | **%OUD** |
|---|---|---|---|
| **Total** | 396 | 478 | 50% |
| **Train** | 308*(77.77%)* | 390*(81.59%)* | 50% |
| **Validation** | 44 *(11.11%)* | 44 *(9.20%)* | 50% |
| **Test** | 44 *(11.11%)* | 44 *(9.20%)* | 50% |

The OUD dataset included 396 participants, comprising 245 males and 151 females. Participant ages ranged from 16 to 79 years, with a mean age of 31 years.
